# The transcriptional regulator Id2 is critical for adipose-resident regulatory T cell differentiation, survival and function

**DOI:** 10.1101/589994

**Authors:** Adolfo B. Frias, Eric J. Hyzny, Heather M. Buechel, Lisa Beppu, Bingxian Xie, Michael J. Jurczak, Louise M. D’Cruz

## Abstract

Adipose regulatory T cells (aTregs) have emerged as critical cells for the control of local and systemic inflammation. In this study, we show a distinctive role for the transcriptional regulator Id2 in the differentiation, survival and function of aTregs. Id2 was highly expressed in aTregs compared with high Id3 expression in lymphoid Tregs. Treg-specific deletion of Id2 resulted in a substantial decrease in aTregs, while Tregs in the spleen and lymph nodes were unaffected. Additionally, loss of Id2 resulted in decreased expression of aTreg associated markers including ST2, CCR2, KLRG1 and GATA3. Gene expression analysis revealed that Id2 expression was essential for the survival of aTregs and loss of Id2 increased cell death in aTregs due to increased Fas expression. Id2-mediated aTreg depletion resulted in increased systemic inflammation, increased inflammatory macrophages and CD8^+^ effector T cells and loss of glucose tolerance under standard diet conditions. Thus, we reveal an unexpected and novel function for Id2 in mediating differentiation, survival and function of adipose-resident Tregs, that when lost resulting in increased metabolic perturbation.

## INTRODUCTION

Visceral adipose tissue (VAT) is enriched for a population of CD4^+^ Foxp3^+^ CD25^+^ Tregs, relative to Treg populations in lymphoid tissue, and that these cells are reduced in frequency and cell number in different models of insulin resistance, including HFD-induced obesity (1). Adipose Tregs (aTregs) are thymically derived (2), maturing in the spleen before seeding the visceral adipose tissue at approximately 10 weeks of age, reaching a peak of ∼40% total CD4^+^ T cells in adipose tissue by ∼25 weeks of age (1, 3). aTreg peripheral development is complex and a recent report indicated a two-step process by which differentiation of these cells occurs (3). aTreg precursor cells can be identified in the spleen by low expression of PPARγ and downregulated expression of the transcriptional regulator Id3 (3, 4). Their TCR repertoire is more restricted than conventional Tregs, suggesting VAT antigen-specific activation followed by clonal expansion (2). A number of surface markers and transcription factors are characteristic of aTregs including ST2, KLRG1, CCR2 and GATA3, and a key feature of these cells is their ∼10-fold increased production of IL-10 relative to conventional Tregs (1, 5, 6).

Detailed gene expression analysis of these cells revealed the transcription factor PPARγ to be highly expressed by aTregs (5, 7). Treg-specific loss of PPARγ did not affect conventional lymphoid-resident Tregs but resulted in a substantial reduction in the frequency and absolute number of aTregs. Moreover, PPARγ-deficient aTregs displayed a gene expression profile more similar to conventional lymphoid Tregs with reduced Foxp3 expression (5). The transcriptional regulators BATF and IRF4 to regulate expression of ST2 and PPARγ in aTregs and IL-33, acting through ST2, is required to both recruit and maintain the aTreg population in adipose tissue (2, 6, 8). Importantly, aTregs are critical for suppressing chronic systemic inflammation observed under conditions of obesity (1, 8). Under HFD conditions aTregs are reduced but can be restored through administration of IL-33 or the thiazolidinedione drug pioglitazone, an approved type II diabetes drug treatment (2, 5, 8). Thus, it is becoming increasingly apparent that aTregs are critical for regulating and maintaining metabolic homeostasis.

The E and Id protein families of transcriptional regulators have been implicated in many aspects of lymphocyte differentiation (9-12). E proteins (e.g. E2A, HEB) are basic helix-loop-helix (HLH) transcriptional activators/repressors that bind to DNA at E-box sites, controlling expression of genes crucial to lineage commitment, targeting antigen-receptor gene rearrangement, and enforcing key developmental checkpoints (9-12). Id proteins (Id1-Id4) heterodimerize with the E proteins, preventing DNA binding at their target E-box site. Thus, Id proteins function as natural dominant-negative regulators of E protein activity.

To investigate the function of E proteins in the thymic development of conventional Tregs, E2A and HEB conditional mice were crossed to a tamoxifen inducible Rosa26-ER-Cre line (13). Administration of tamoxifen induced deletion of the E proteins E2A and HEB and resulted in a dramatic increase in Foxp3^+^ Tregs in the thymus (13). Moreover, conditional loss of Id2 and Id3 using CD4-Cre led to a decrease in thymic Foxp3^+^ Tregs (13). These data strongly suggest a role for the E and Id proteins in the regulation of lymphoid Tregs. However, the use of non-Treg specific Cre lines and a focus on analysis of the thymically-derived Treg populations, precluded significant advancement in understanding the function of these proteins in aTregs. More recently, E2A and HEB were shown to be critical for effector Tregs with Foxp3-Cre mediated loss of E proteins resulting in substantial increases in Tregs in peripheral tissues such as the small intestine, lung and liver, although the Treg population residing in adipose tissue was not examined (14). The function of Id2 and Id3 in conventional Tregs has also been examined by crossing Id2 and Id3 floxed animals to a Foxp3-Cre line (15). Treg-specific loss of both Id2 and Id3 resulted in a decrease in Tregs in lymphoid tissues and a corresponding increase in spontaneous inflammation and autoimmunity in these animals (15). Another study analyzed the function of Id3 in germline deficient mice, concluding loss of Id3 resulted in the generation of fewer Tregs in the thymus and spleen (16). However, the function of the Id proteins in aTregs was not examined in either of these studies. Using Id3-GFP reporter mice, it was recently shown that Id3 expression is gradually reduced in effector regulatory T cells as they seed peripheral tissue (4). Therefore, although it is clear that E and Id proteins play reciprocal roles in thymic Treg development and homeostasis, the function of these proteins in aTregs has not been determined.

Here we examined the role of Id2 in the differentiation, survival and function of aTregs. We show that Id2 is highly upregulated in aTregs, while Id3 is expressed predominantly in lymphoid-resident Tregs. Id2 was required for the survival of aTregs and its loss resulted in substantially decreased aTreg population. Moreover, RNA-seq analysis revealed increased Fas expression in Id2-deficient aTregs, providing a possible mechanistic explanation for the increased cell death we observed in the absence of Id2. Physiologically, loss of Id2 in Tregs resulted in increased systemic inflammation, inflammatory macrophage and CD8^+^ effector T cell infiltration and glucose intolerance under standard diet conditions, indicating Id2 was critical for the survival and function of aTregs in order to maintain adipose-tissue homeostasis.

## MATERIALS AND METHODS

### Mice

All experiments were approved by the University of Pittsburgh Institutional Animal Care and Use Committee. The Foxp3YFPiCre mice were obtained from Jackson Laboratories (17). The Id2 floxed mice were a gift from Anna Lasorella (Columbia University), Id2-YFP mice from Ananda Goldrath (UC San Diego), and Id3-GFP mice from Cornelis Murre (UC San Diego). Mice were bred and housed in specific pathogen-free conditions in accordance with the Institutional Animal Care and Use Guidelines of the University of Pittsburgh.

Mixed bone marrow chimeras were generated by transferring either 5 × 10^6^ B220^-^CD3^-^ NK1.1^-^ bone marrow cells from a CD45.2 Foxp3YFPiCre^+^ donor WT donor or 5 × 10^6^ B220^-^ CD3^-^NK1.1^-^ bone marrow cells from a CD45.2 Id2^f/f^ Foxp3YFPiCre^+^ donor into lethally irradiated (1000 rad) CD45.1 recipient mice. All chimeras were rested at least 8 weeks to allow reconstitution of the host.

For the HFD studies, mice were fed a rodent diet of 60% kcal fat from Research Diets (D12451) for 12 weeks.

### Flow cytometry and cytokine stimulation assay

Single-cell suspensions were prepared from thymus and spleen. Bone marrow was prepared by isolating lymphocytes from the femurs of mice and preparing a single-cell suspension. Gonadal adipose single-cell suspensions were prepared as previously described (18). All antibodies were purchased from BD Biosciences and eBioscience (San Diego, CA, USA) unless otherwise specified. Samples were collected on a Cytek Aurora (Cytek Biosciences), FACS LSRII, FACS Fortessa or FACS Aria (BD Biosciences, San Jose, CA, USA) and were analyzed with Flowjo software (TreeStar, Ashland, OR, USA).

For intracellular cytokine staining we stimulated the cells with PMA (50ng/ml) (Sigma) and ionomycin (1nM) (Sigma) for 4 hours. We added protein transport inhibitor (eBiosciences) to the culture at the recommended concentration. Cells were surfaced stained, before fixing, permeabilizing and intracellular staining according to the manufacturer’s instructions (eBioscience Foxp3 staining kit).

### Quantitative PCR

Total RNA was isolated from spleen or adipose tissue single cell suspensions using the RNeasy Mini Purification Kit (Qiagen) and cDNA was obtained using Superscript First Strand cDNA synthesis kit (GeneCopoeia). qPCR was performed using Taqman probes for the indicated gene (Applied Biosystems). Target gene expression was normalized to housekeeping gene *HPRT*.

### RNA sequencing and analysis

RNA was isolated from sorted Tregs (CD4^+^, CD25^+^, CD127^-^) from the spleen or adipose tissue. Libraries were prepared using Ultra DNA library preparation kits. RNA sequencing analysis was performed on Illumina NextSeq500 using 500bp paired-end reads by Health Science Sequencing core facility at University of Pittsburgh. Raw sequence reads were trimmed of adapter sequences using CLC Genomics Suite. The trimmed reads were mapped onto the mouse genome and gene expression values (TPM; transcripts per million kilobase) were calculated using CLC Genomics Suite Workbench 11. Differential gene expression was analyzed using Partek Genomics Suite and graphs generated using Graphpad Prism with values normalized as follows: (TPM value for gene X in condition A)/ (mean TPM of gene X in all conditions for that sample).

### Determination of fasting blood glucose, insulin and glucose tolerance testing

Mice were fasted overnight before fasting blood and insulin levels were measured. Blood glucose was measured using a handheld glucometer (Contour NEXT EZ, Bayer) and plasma insulin was measured by ELISA (Alpco). For the glucose tolerance tests (GTTs), we administered glucose (2g/kg of body weight) by intraperitoneal injection after an overnight fast. We measured changes in plasma insulin at 15, 30 and 120 minutes and blood glucose at 15, 30, 45, 60 and 120 minutes after glucose injection.

### Statistical analysis

All graphs were created using GraphPad Prism 7 and statistical significance was determined with the two-tailed unpaired Student’s t-test or using one-way ANOVA adjusted for multiple comparisons where appropriate.

## RESULTS

### Id2 and Id3 are differentially expressed in aTregs

We first determined the expression of Id2 and Id3 mRNA in aTregs. Sorting CD4^+^ CD25^+^ Tregs from the spleen and visceral adipose tissue (VAT) of 25-week-old male C57BL/6 mice, we found that Id2 mRNA was expressed at ∼4-fold higher in aTregs relative to splenic Tregs (Fig. 1A). In contrast, at the mRNA level, Id3 expression appeared approximately equal in splenic and aTregs (Fig. 1A). To confirm these findings, we used Id2-YFP and Id3-GFP reporter mice. In agreement with our qPCR data, gating on Tregs from the spleen or VAT, we determined high Id2-YFP expression in aTregs compared to their splenic counterparts (Fig. 1B). We also determined low Id3-GFP expression in aTregs and high expression in splenic Tregs (Fig. 1B). These data are consistent with recently published data determining that Id3 expression is high in splenic Tregs and reduced in tissue Tregs (4). Additionally, gating on Id2-YFP^+^ and Id2-YFP^-^ Tregs in the spleen, we noted higher KLRG1 expression in the Id2-YFP^+^ Tregs and slightly elevated ST2 expression, while the converse was true of Id3-GFP^+^ versus Id3-GFP^-^ Tregs (Fig. 1C). KLRG1 is a marker associated with effector Tregs while ST2, the IL-33 receptor, is upregulated after Tregs migrate to peripheral tissue (3). Therefore, these data support a model whereby loss of Id3 and concurrent upregulation of Id2 marks effector-like Tregs that seed peripheral tissue, including the VAT (3, 4). We concluded from these data that Id2 is preferentially expressed by aTregs, while Id3 is expressed by lymphoid tissue resident Tregs.

**Figure 1.**
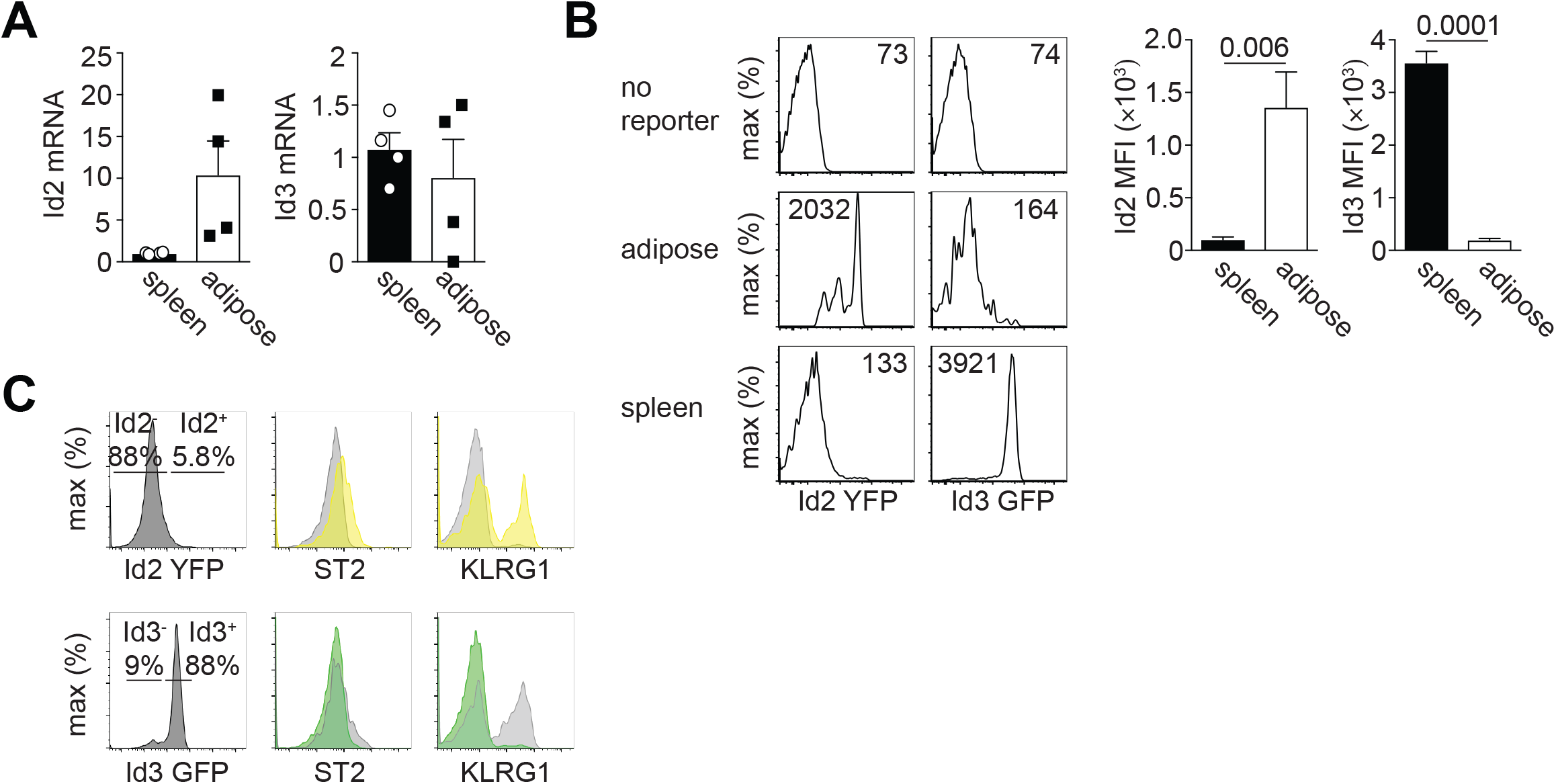
Id2 and Id3 expression in aTregs. (A) qPCR showing Id2 and Id3 mRNA expression in sorted CD4^+^ CD25^+^ Tregs isolated from the spleen or visceral adipose tissue. Id2 or Id3 expression was normalized to HPRT. Each dot indicates cells isolated from one animal. (B) Flow cytometry histograms showing Id2-YFP or ID3-GFP expression in gated CD4^+^ CD25^+^ Tregs isolated from the spleen or visceral adipose tissue. B6 mice were used as a ‘no reporter’ control. Numbers in histograms indicate the median fluorescence intensity (MFI). Bar graphs indicate the average MFI in Tregs isolated from the indicated tissue. (C) Histograms indicating the frequency of Id2^+^ or Id3^+^ Tregs identified in the spleen (left) and their expression by flow cytometry of ST2 and KLRG1 (right). Yellow peaks indicate Id2-YFP^+^ cells and green peaks indicate Id3-GFP^+^ cells. Grey peaks indicate Id2-YFP^-^ or Id3-GFP^-^ cells, respectively. Data are representative of two independent experiments with 1-3 mice per group. P values were calculated using the student’s t-test.

### Id2 is critical for maintenance of aTregs

To determine the specific function of Id2 in Tregs, we crossed Id2 floxed mice to the Foxp3-YFP-iCre line (17) to generate mice in which Id2 was conditionally deleted in Tregs. We called these mice Id2 CKO (conditional knock out) and compared them to Foxp3-YFP-iCre only littermate controls (Ctrl, WT). We first examined the frequency of Tregs in adipose tissue, lymph nodes (LNs) and spleen in the absence of Id2 expression. Gating on total CD4^+^ T cells, we observed that Treg-specific loss of Id2 expression resulted in a significant reduction of Foxp3^+^ CD25^+^ aTregs but not Tregs in the LNs and spleen (Fig. 2A). The frequency of Tregs in the thymus was not affected by the absence of Id2 (data not shown).

**Figure 2.**
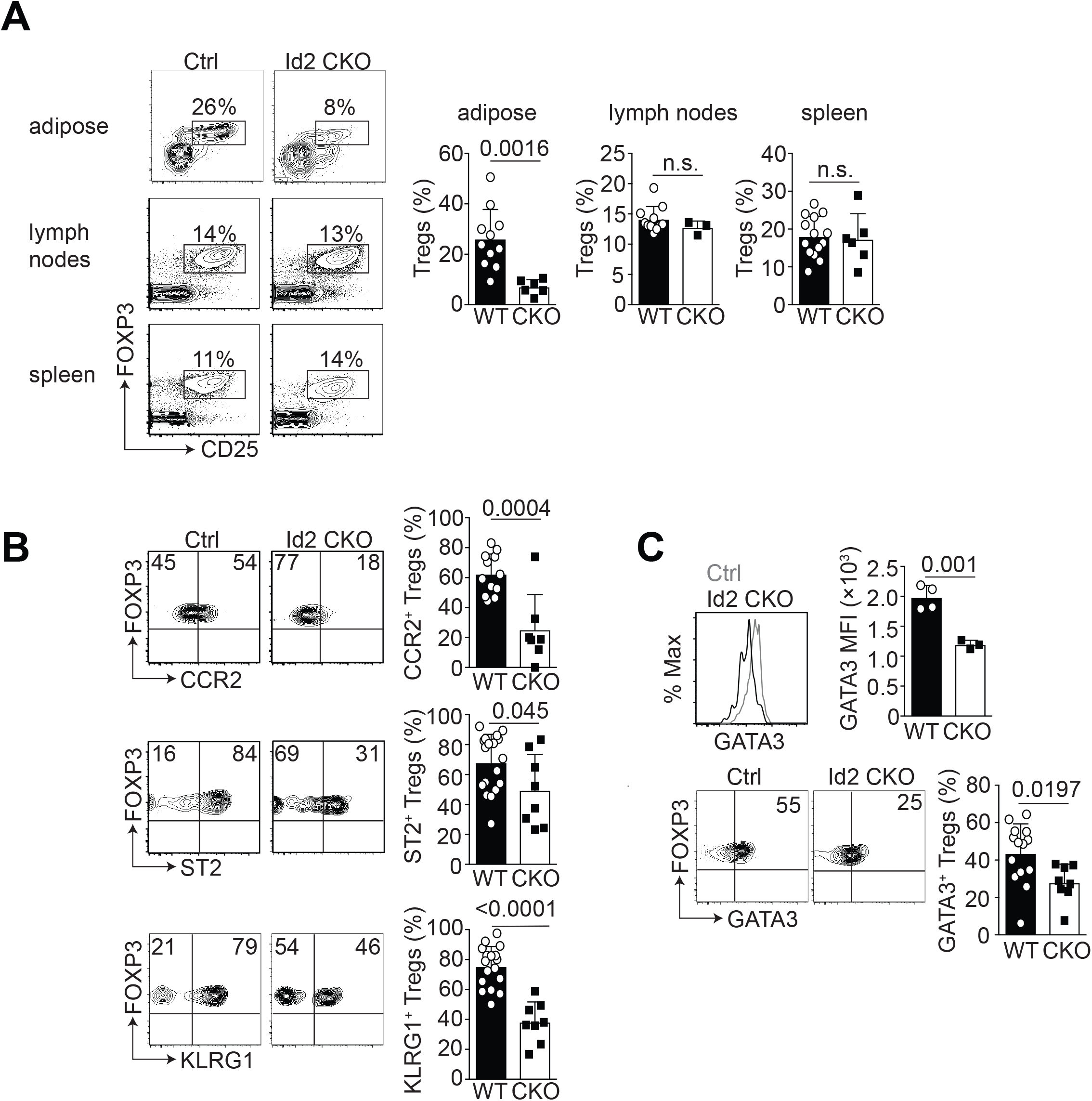
Id2-deficiency results in reduced frequency of aTregs. (A) Flow cytometry plots showing the frequency of Foxp3^+^ CD25^+^ gated CD4^+^ T cells from the indicated tissue in Control (Ctrl) and Id2-deficient (Id2 CKO) mice. Bar graphs indicate the frequency of Foxp3^+^ CD25^+^ Tregs from the adipose, lymph nodes or spleen. Each dot indicates one animal. (B) Flow cytometry plots showing CCR2, ST2 and KLRG1 expression on gated Foxp3^+^ CD25^+^ CD4^+^ T cells in the adipose tissue in Ctrl or Id2 CKO mice. Bar graphs indicate the frequency of CCR2^+^, ST2^+^ and KLRG1^+^ aTregs from wildtype (WT) and Id2 CKO (CKO) mice. (C) Histogram, flow cytometry plots and bar graphs indicating the median fluorescence intensity (MFI) and frequency of GATA3 expression in gated aTregs from WT and Id2 CKO mice. Data are representative of at least four independent experiments with 1-5 mice per group. P values were calculated using the student’s t-test.

We next assessed the phenotype of the aTreg, focusing on the markers CCR2, ST2 and KLRG1, comparing the aTreg in WT and Id2-deficient mice. Expression of CCR2, ST2 and KLRG1 were all significantly reduced in the Id2-deficient aTregs relative to the WT (Fig. 2B). We also determined expression of GATA3, a transcription factor associated with tissue-resident Tregs. Notably, GATA3 expression was reduced in Id2-deficient aTregs relative to WT Tregs (Fig. 2C). We also examined expression of Foxp3 in these cells and observed no difference in Foxp3 expression in the absence of Id2 in aTregs (Supplemental Figure 1). Thus, these data show that Id2 expression is required for the maintenance of the aTregs population and for differentiation of the aTreg phenotype.

### Id2 expression is required for aTreg survival

As we observed a substantial reduction in the frequency of the aTreg population in the absence of Id2 (Fig. 2A), we next wanted to determine the cause of this decrease in aTregs. To address this, we performed transcriptomic analysis of Tregs sorted from the spleen or visceral adipose tissue of >15-week old WT or Id2 CKO male mice. Genes previously reported to be associated with aTregs were substantially upregulated in our sorted aTreg dataset compared to splenic Treg, indicating our gene expression analysis was robust (data not shown). Closer inspection revealed that a number of genes associated with cell death and survival, transcription and metabolism were up-or downregulated in the Id2 CKO aTregs, relative to splenic Tregs (Fig. 3A). In particular, we were intrigued by the upregulation of Fas in the Id2-deficient aTregs as these data suggested that Id2-deficient aTregs may be more prone to cell death.

**Figure 3.**
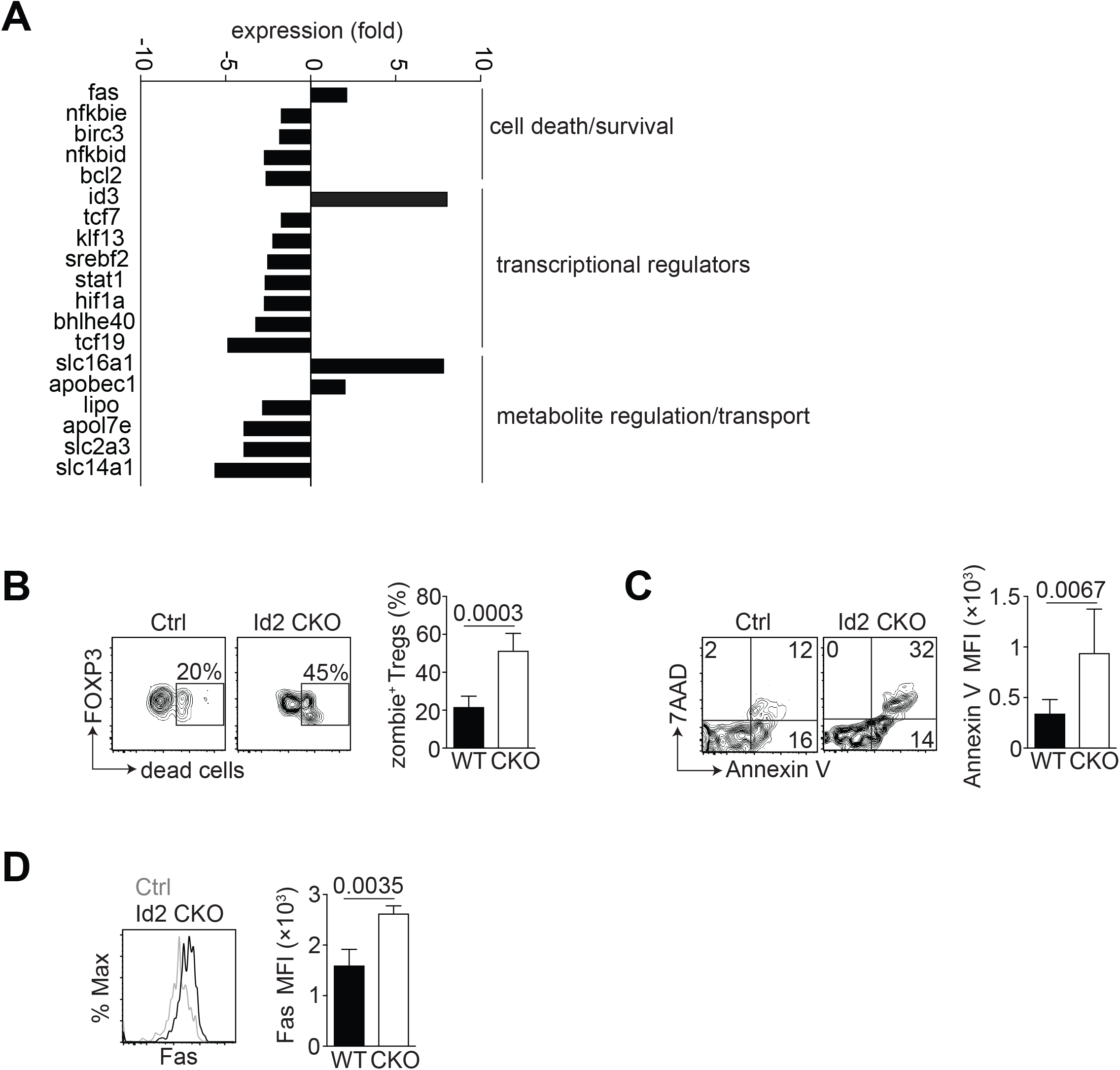
Loss of Id2 decreases survival of aTregs. WT and Id2 CKO aTregs were sorted from the spleen or adipose tissue of 15-20-week-old male mice for RNA-sequencing. (A) Bar graphs indicating expression of transcripts involved in various pathways (indicated) in Id2 CKO aTregs relative to WT aTregs. (B) Flow cytometry plots and bar graph from Ctrl and Id2 CKO mice showing the frequency of viability dye (zombie)^+^ gated CD4^+^ Foxp3^+^ aTregs. (C) Flow cytometry plots and bar graph indicating the Annexin V frequency and MFI on WT vs CKO aTregs. (D) Histogram and bar graph indicating Fas MFI in WT vs CKO aTregs. Data are representative of two independent experiments with 1-3 mice per group. P values were calculate using the student’s t test.

To confirm that Id2 was required for the survival of a Tregs, we used a fixable viability ‘zombie’ dye and noted an increase in dead aTregs in the absence of Id2 relative to WT aTregs (Fig. 3B). Moreover, a greater frequency of Id2-deficient aTregs stained positive for AnnexinV and 7AAD compared to their WT counterparts (Fig. 3C). When we tested aTregs for expression of Fas, we observed Id2-deficient aTregs express more Fas relative to their WT counterparts (Fig. 3D), which was consistent with our gene expression analysis. We thus concluded that one mechanism by which Id2 regulates aTregs is by promoting their survival in VAT.

From our gene expression analysis, we had also noted the decreased expression of *Hif1α* in the Id2-deficient aTregs (Fig. 3A). Hif1α (Hypoxia Inducible Factor 1α) is critical for cell survival under low oxygen conditions. Relative to splenic Tregs, we observed more aTregs express Hif1α (>50%) (Supplemental Figure 2). However, in agreement with our gene expression data, Id2-deficiency resulted in reduced Hif1α protein expression on aTregs. Taken together, our data show that Id2 is critical for expression of proteins required for the survival of aTregs.

### Id2 expression is required for aTreg cytokine production

Previous data have shown aTregs are potent IL-10 producers (1, 6) and it is likely this IL-10 contributes to an anti-inflammatory environment in non-obese adipose tissue. Additionally, Il-13 production by Tregs was recently shown to support macrophage IL-10 secretion and efferocytosis to promote inflammation resolution (19). Stimulation of WT or Id2-deficient aTregs *in vitro* revealed reduced production of IL-10 and IL-13 by the Id2-deficient aTregs (Fig. 4A and B). We could not detect a difference in IFN-γ production by aTregs from WT versus CKO mice (Fig. 4C). Overall, our data suggest that loss of Id2 resulted in reduced anti-inflammatory cytokine production by aTregs.

**Figure 4.**
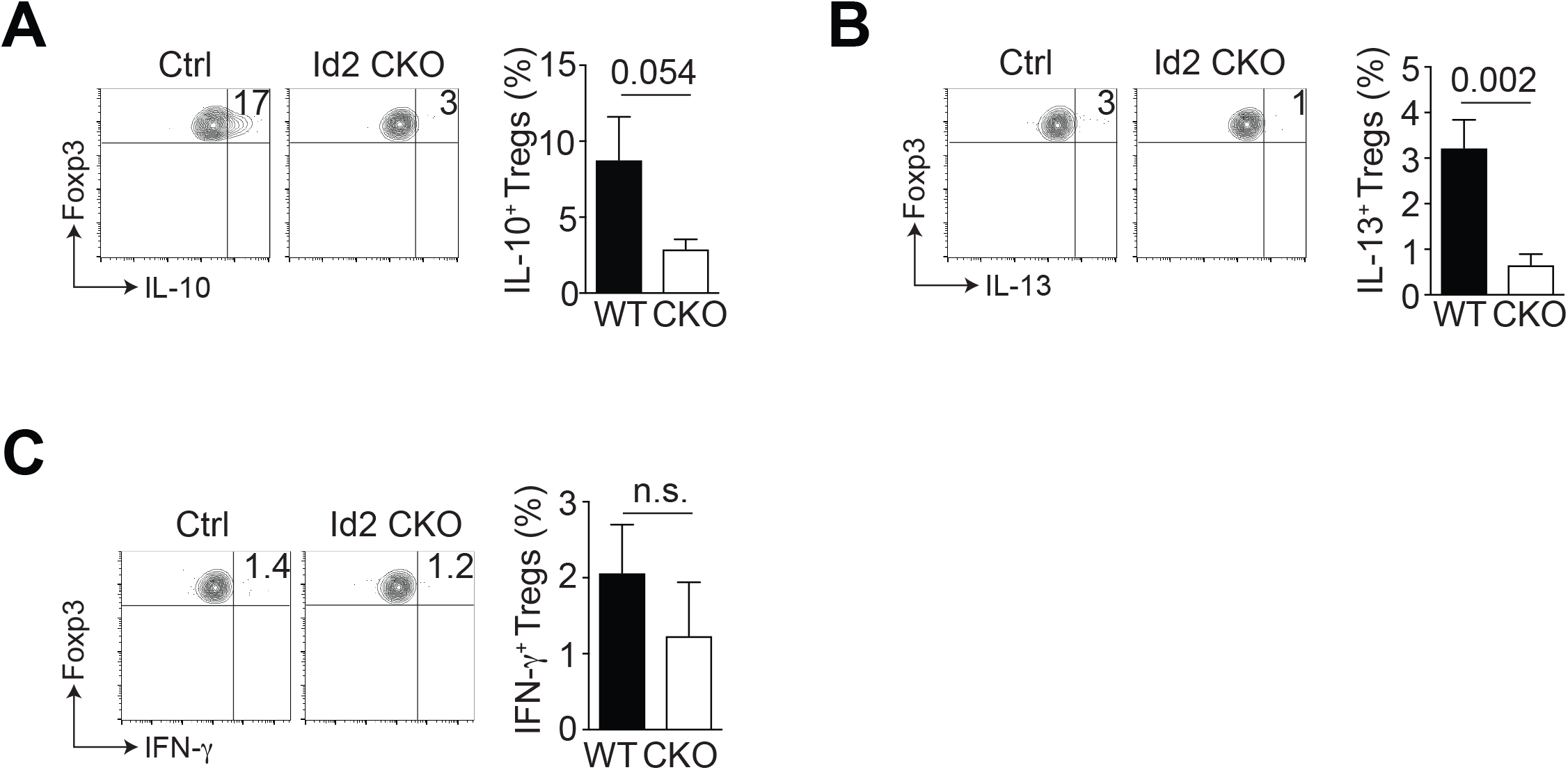
Cytokine production by Id2-deficient aTregs. The total stromal vascular fraction (SVF) isolated from the adipose tissue of WT or Id2 CKO mice was cultured with PMA and ionomycin for 4 hours and cytokine production by gated CD4^+^ Foxp3^+^ aTregs analyzed by flow cytometry. (A) Flow cytometry plots and bar graph indicating IL-10 expression by gated CD4^+^ Foxp3^+^ WT vs CKO aTregs. (B) Flow cytometry plots and bar graph indicating IL-13 expression by gated CD4^+^ Foxp3^+^ WT vs CKO aTregs. (C) Flow cytometry plots and bar graph indicating IFN-γ expression by gated CD4^+^ Foxp3^+^ WT vs CKO aTregs. Data are representative of two independent experiments with 2-3 mice per group. P values were calculate using the student’s t test.

### Loss of Id2 in aTregs results in reduced aTreg frequency under standard or high fat diet conditions

It has previously been reported that under high fat diet (HFD) conditions, there is a loss of Tregs from the VAT (1, 5, 6). To test this, and whether the requirement for Id2 in Tregs was cell intrinsic, we generated bone marrow chimeras (BMC) in which we transferred congenically marked WT or Id2 CKO bone marrow into sublethally irradiated CD45.1 hosts. At four weeks post-reconstitution, we placed these chimeric mice on high fat diet (HFD) for 12 weeks, leaving the control groups on control diet (CD). As has been reported previously, there was a significant reduction in the frequency of aTregs in WT mice on HFD relative to their CD counterparts (Fig. 5A). Importantly, we observed that the reduction in aTregs in mice with loss of Id2 expression on CD was more severe than in WT mice on HFD (Fig. 5A) (1). Additionally, ST2, CCR2, KLRG1 and GATA3 were all reduced in the Id2-deficient aTregs, irrespective of their diet (Fig. 5A). We concluded that Treg-specific deficiency in Id2 resulted in loss of aTregs that was comparable with the frequency of aTregs in WT mice on HFD.

**Figure 5.**
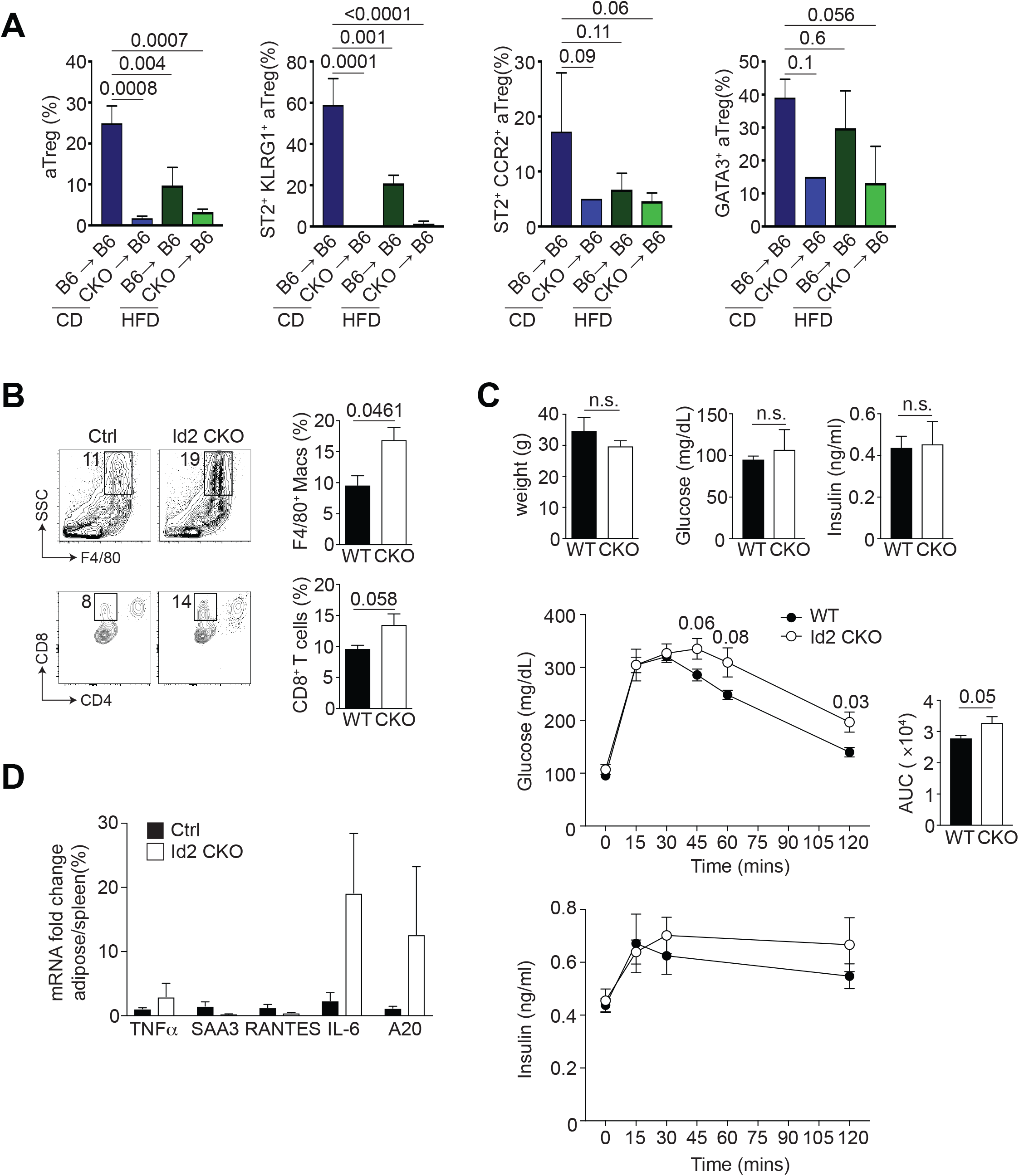
Physiological function of Id2^+^ aTregs. (A) CD45.1 mice were irradiated and injected with bone marrow cells from WT or Id2 CKO mice. Four weeks after reconstitution, mice were placed on CD or HFD for 12 weeks. Bar graphs indicating the frequency of CD4^+^ Foxp3^+^ aTregs, ST2^+^ KLRG1^+^, ST2^+^ CCR2^+^ and GATA3^+^ gated aTregs in the indicated BMC under CD or HFD conditions. (B) Flow cytometry and bar graphs indicating the frequency of total macrophages (F4/80^+^), M1 (CD11c^+^ F4/80^+^), M2 (CD206^+^ F4/80^+^), IL-10^+^ F4/80^+^ macrophages and CD8^+^ T cells isolated from the adipose tissue of WT or Id2 CKO mice on CD. (C) Graphs indicating weight, fasting glucose, fasting insulin and blood glucose and insulin over time following GTT. Calculated area under the curve (AUC) from all mice tested by GTT. (D) Graph indicating the expression of TNFα, SAA3, RANTES, IL-6 and A20 mRNA fold change in total adipose tissue SVF versus spleen. Data are representative of two independent experiments with 2-3 mice per group (A, B and D) and one experiment with 6 mice per group (C). P values were calculate using the student’s t test or one-way ANOVA.

### Id2-deficiency in aTregs results in perturbed systemic inflammation and metabolism

Loss of aTregs due to genetic perturbations or HFD can also affect other adipose-resident immune cells (1, 5, 6). We tested how loss of Id2 in aTregs affected other adipose-resident cells and found a significant increase in F4/80^+^ macrophages in the adipose tissue of Id2 CKO mice (Fig. 5B). Moreover, the frequency of CD11c^+^ F4/80^+^ M1 macrophages was substantially increased in the absence of Id2, although the CD206^+^ M2 macrophages remained unchanged (Fig. 5B). When we cultured the total stromal vascular fraction (SVF) from the adipose tissue with PMA and ionomycin, we noted that IL-10 production by the F4/80^+^ macrophages was substantially reduced in the Id2 CKO cohort (Fig. 5B). Additionally, we observed an increased frequency of CD8^+^ effector T cells in the absence of Treg-specific Id2 (Fig. 5B). Taken together, these data show that there is an increase in pro-inflammatory macrophages and CD8^+^ effector T cells in the absence of Id2^+^ aTregs.

To determine how Id2-mediated aTreg loss affected glucose metabolism and insulin action in these animals, we fasted CD WT and Id2 CKO mice overnight and measured their fasting blood glucose and plasma insulin levels. Although weight, fasting glucose and insulin levels were unaffected by Id2-specific loss in Tregs, Id2 CKO mice displayed impaired glucose tolerance during the GTT (Fig. 5C). Plasma insulin levels were also slightly elevated in the glucose-treated Id2 CKO mice over time, relative to the WT mice (Fig. 5C). Additionally, mRNA analysis revealed that inflammatory mediators such as *Tnfα, Il6* and *A2*0 (*Tnfaip*3) were all increased in total adipose tissue relative to spleen in the Id2-deficient animals (Fig. 5D). Together our data show that loss of Id2 in Tregs in mice on CD results in increased inflammation, increased inflammatory immune cells and metabolic perturbation comparable with WT mice on HFD.

## DISCUSSION

The role of the immune system in influencing systemic chronic inflammation, insulin sensitivity and glucose tolerance associated with obesity and adipose tissue has recently been elucidated. We now know that increasing adiposity resulting from increased caloric intake results in reduction in immune cells associated with anti-inflammation such as Th2 cells, M2 macrophages, ILC2s and aTregs and a corresponding increase in inflammatory cells such as M1 macrophages, Th1 cells and cytotoxic CD8^+^ T cells (20). In our work here, we explore the transcriptional requirements of aTregs and show the transcriptional regulator Id2 is critical for their survival and function.

In other immune cell contexts, it was previously shown that Id2 is critical for cell survival. In the CD8^+^ T cell immune response to infection, Id2 is substantially upregulated during the effector response and loss of Id2 resulted in loss of responding effector T cells due to unrestricted E protein activity (21, 22). Similarly, in invariant Natural Killer T (iNKT) cells, Id2 was critical for survival of hepatic iNKT cells and loss of Id2 resulted in loss of Bcl-2 expression and increased cell death of these cells (23). In peripheral Tregs, recent gene expression analysis revealed loss of E proteins resulted in upregulation of proteins associated with proliferation and survival such as Bcl-2, Ki-67 and Bcl-11b (14). Here we showed that Id2 is a critical factor for the survival of aTregs and that loss of Id2 results in upregulation of the pro-apoptotic protein Fas. Notably, we did not see upregulation of Bim expression in the Id2-deficient aTregs (data not shown). However, we did observe that Id3 was substantially upregulated in the absence of Id2, suggesting that Id3 upregulation can compensate for some Id2 loss and negatively regulate E protein transcriptional activity at certain genomic sites.

One interesting chemokine associated with aTregs is CCR2, the receptor for CCL2 which is predominantly expressed on macrophages and monocytes, but is also associated with activated T cells (24). With loss of Id2 in aTregs, we observed reduced CCR2 expression, leading to concerns that migration of aTregs from the spleen to the adipose tissue affected their accumulation in the adipose tissue. However, loss of CCR2 on T cells resulted in increased accumulation and function of Tregs in the spleen and colon (24), suggesting that reduced CCR2 expression in the absence of Id2 should not affect migration of aTregs.

Regarding function of Id2-deficient aTregs, we observed increased localized adipose tissue inflammation, increased inflammatory macrophage and CD8^+^ T cell infiltration and impaired glucose tolerance after fasting in the Id2 Treg-deficient animals relative to WT controls. Additionally, we detected reduced IL-10 and IL-13 production by the Id2-deficient aTregs upon stimulation *in vitro*. IL-10 has been shown to be positively regulated by E proteins in conventional T cells (25). However, in peripheral tissue Tregs it was recently reported that IL-10 is likely negatively regulated by E2A and HEB (14), suggesting that E protein activity in T cells is context-dependent. With loss of Id2 and unchecked E protein activity, we see reduced IL-10 production, suggesting that 1) IL-10 is negatively regulated by E2A and HEB in aTregs and 2) loss of Id2 in aTregs reduces their ability to suppress local and systemic inflammation, resulting in perturbed glucose tolerance.

Our finding that Hif1α is highly expressed in aTregs and reduced in the absence of Id2 suggests a previously unappreciated mechanism for aTreg maintenance and survival. Hif1α has previously been described as upregulated in adipocytes under HFD conditions (26, 27), presumably to influence survival under increasing hypoxic conditions, and its expression is also associated with induction of Foxp3, increasing abundance and function of Tregs (28). We propose here that loss of Hif1α in the absence of Id2 in aTregs helps to contribute to their increased cell death and loss of functionality in adipose tissue, thus providing new insights into the transcriptional regulation and survival of adipose-resident Tregs.

In summary, here we provide evidence of a new role for the transcriptional regulator Id2 in the homeostasis, differentiation, survival and function of aTregs. Loss of Treg-specific Id2 did not impact Tregs in the lymphoid tissues but led to dramatic loss of these cells in adipose tissue. We showed that Id2 expression was critical for the survival of aTregs and that loss of these cells resulted in increased local and systemic inflammation, increased immune infiltration into adipose tissue and impaired glucose tolerance.

## AUTHORS CONTRIBUTIONS

A.B.F. and E.J.H. conducted the experiments and wrote the paper; H.M.B. and L.B. conducted the experiments; B.X. and M.J.J. performed the metabolic studies; L.M.D. designed the experiments and wrote the paper.

## ACKNOWLEDGMENTS

We would like to acknowledge Drs. McGeachy and Kane for critical reading the manuscript and all members of the D’Cruz lab for their constructive criticism and comments. We would also like to thank the University of Pittsburgh Unified Flow Core for assistance with cell sorting and flow cytometry. The authors declare no competing interests. This work was supported by NIH R01 DK114012 and R01 DK119627 to M.J.J. and seed funding from the University of Pittsburgh to L.M.D.

