## Supplemental Figure 1 for "The transcriptional regulator Id2 is critical for adipose-resident regulatory T cell differentiation, survival and function"

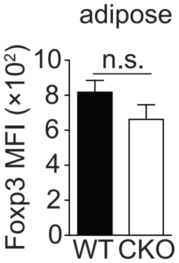


**Supplemental Figure 1. Foxp3 MFI is unchanged in Id2-deficient aTregs.** Bar graph indicating the median fluorescent intensity (MFI) of Foxp3 in WT or Id2 CKO Tregs isolated from the adipose tissue. Data are representative of three independent experiments.
