## Supplemental figure 2 for "The transcriptional regulator Id2 is critical for adipose-resident regulatory T cell differentiation, survival and function"

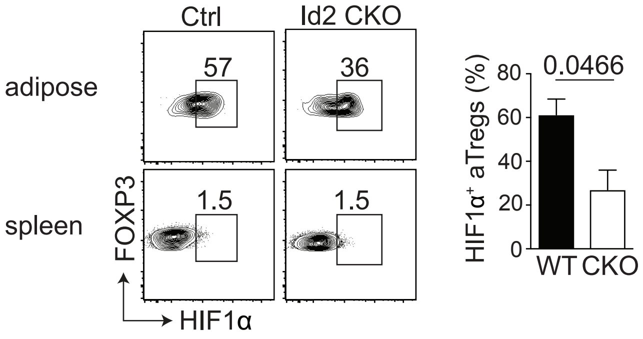


**Supplemental Figure 2. Hif1α expression in Id2-deficient aTregs.** Flow cytometry plots and bar graph indicating the frequency of Hif1α+ Foxp3+ aTregs from WT and Id2 Treg-specific deficient male mice. Data are representative of two experiments with 2 mice per group. P values were calculate using the student's t test.
